# Spinach Growers Value Host Resistance and Synthetic Fungicides in the Fight against Downy Mildew

**DOI:** 10.1101/2020.01.20.913129

**Authors:** Robin A. Choudhury, Neil McRoberts

## Abstract

California spinach growers struggle to manage spinach downy mildew disease. The disease is especially difficult in the organic crop, which currently relies on resistant varieties to maintain disease-free crop. Alternative control measures are available, but it is not clear how growers perceive the efficacy of these methods. It is also not clear who growers contact to find out information on spinach downy mildew disease management. In this study, we conducted an online survey of people involved in spinach production, asking about their beliefs in the efficacy of different control methods and who they contact frequently to discuss spinach downy mildew control. We found that respondents were most positive about the efficacy of resistant varieties and synthetic pesticides, with much lower perceived efficacy for the practices of disking diseased fields, roguing diseased plants, and organic pesticides. Growers most frequently contacted pest control advisors (PCAs) about management strategies for spinach downy mildew. These results suggest that respondents are most confident about the efficacy of resistant varieties and synthetic pesticides and may be hesitant to adopt new control strategies like organic pesticides. The results also suggest that future extension efforts can be focused on PCAs to reach the most stakeholders with up to date research on downy mildew control.

## Introduction

Spinach is a high value crop that California leads the nation in production. In California, baby leaf spinach production is typically produced on 80-inch beds, allowing for dense plantings and more consistent growth. Baby leaf spinach is typically harvested within approximately 30-45 days after planting, allowing for high throughput on land production. Some producers in the Salinas Valley still produce freezer spinach, which can be in the ground for 90-120 days and is harvested several times (Choudhury et al. 2016a; Koike et al. 2011). Many spinach producers harvest their spinach multiple times by cutting the crop and re-sprouting the unharvested portions.

The most important constraint for spinach producers in California is spinach downy mildew, caused by the oomycete *Peronospora effusa* (syn. *P. farinosa spinacae*) (Choudhury et al. 2016a; Kandel et al. 2019). Spinach downy mildew is an obligate biotrophic pathogen and requires a living host to grow and reproduce. Spinach downy mildew causes chlorotic lesions on the leaves, where even low levels of disease (e.g. 1% disease incidence) cannot be accepted for sale to consumers (Choudhury et al. 2016a). Spinach downy mildew is especially constraining on organic production, which cannot rely on synthetic fungicides to help control the disease. Most organic growers rely heavily on disease resistant varieties. As new races of the pathogen overcome the resistant varieties, growers need to heavily rely on new varieties being bred by a relatively small portion of seed producers (Feng et al. 2018). Although some organic fungicides on the market are available and show some control of disease, they do not control to economic levels and need to be supplemented with resistant varieties for full effect (Choudhury et al. 2015a; b; Choudhury et al. 2017b).

Baby spinach production in California is a complex affair, requiring many different groups cooperating and working together to produce healthy crops. Growers typically operate in groups, with field managers directing crews of workers who help to plant, maintain and harvest the crop. Growers typically work closely with packers and shippers, who contract with growers for spinach over the course of the year.

Seed breeders and producers grow seeds in cool climates that have long daylengths to promote bolting and seeding of spinach, such as the Netherlands and Washington state. Pest control advisors (PCAs) work closely with growers and are responsible for monitoring the crop for any disease or pest issues and make recommendations on what products to spray to help maintain a healthy crop. Public researchers investigate different aspects of spinach production and share their research openly. Extension agents conduct field trials testing out practical control strategies, often in close collaboration with growers and PCAs, and also help to make recommendations based on the latest research from public researchers.

## Survey Instrument Design

We focused our survey design on a handful of questions to minimize the length of the survey. We sent our survey by email to 210 people involved with the California Leafy Greens Research Board. There were nine survey questions that focused on three main concerns: the role of the participant in spinach production and who else within the spinach production community they communicate with about spinach downy mildew control, the perceived efficacy of different management practices and frequency of disease, and the demographics of the participants. The survey conducted online between 2017-06-07 and 2017-07-31, and 53 people responded to the survey. Survey questions were designed to fill in gaps of knowledge based on in-field discussions with growers and PCAs about management practices and communication. Our survey was certified by the UC Davis Institutional Review Board (IRB identification number 935570-1).

## Communication About Spinach Downy Mildew Management

Survey participants were asked what group they most closely associated with and how frequently they contacted other groups about spinach downy mildew information. Participants that described their group as ‘other’ were categorized to the nearest relevant production group. Frequency of contact responses were coded as zero through four, and the median value was used for interpretation of results.

Pest control advisors and packers and shippers frequently contacted all other groups about spinach downy mildew information. The most contacted groups were growers and breeders and seed producers. Public researchers rarely contacted others within production for information about spinach downy mildew, including extension.

It is possible that these results were limited by the forced groupings of participants, where some participants may identify equally with multiple groups (e.g. – extension and public research) but for the analyses they could only choose one role. The survey sample size was also relatively small, with only 53 respondents. The online survey format provided a fast and easy way to contact many people involved in production, however several studies have found that in-person surveys have higher participation rates and (in some cases) fewer artifacts of the research.

## Perceived Efficacy of Different Disease Management Practices

Many respondents perceived that resistant seed varieties and synthetic pesticides were the most efficacious treatments for spinach downy mildew. These management strategies are currently considered the standard for most conventional growers. Resistant seed varieties provide qualitative resistance, allowing growers to produce large plots of healthy plants. However, the longevity of resistant genes is relatively short, as new races of spinach downy mildew rapidly emerge to overcome host resistances. In addition, the high costs associated with seeds of resistant varieties is prohibitive to some growers. Although most *P. effusa* populations are susceptible to synthetic pesticides, these treatment types are unavailable for organic producers, limiting their use for widespread control.

**Figure 1:**
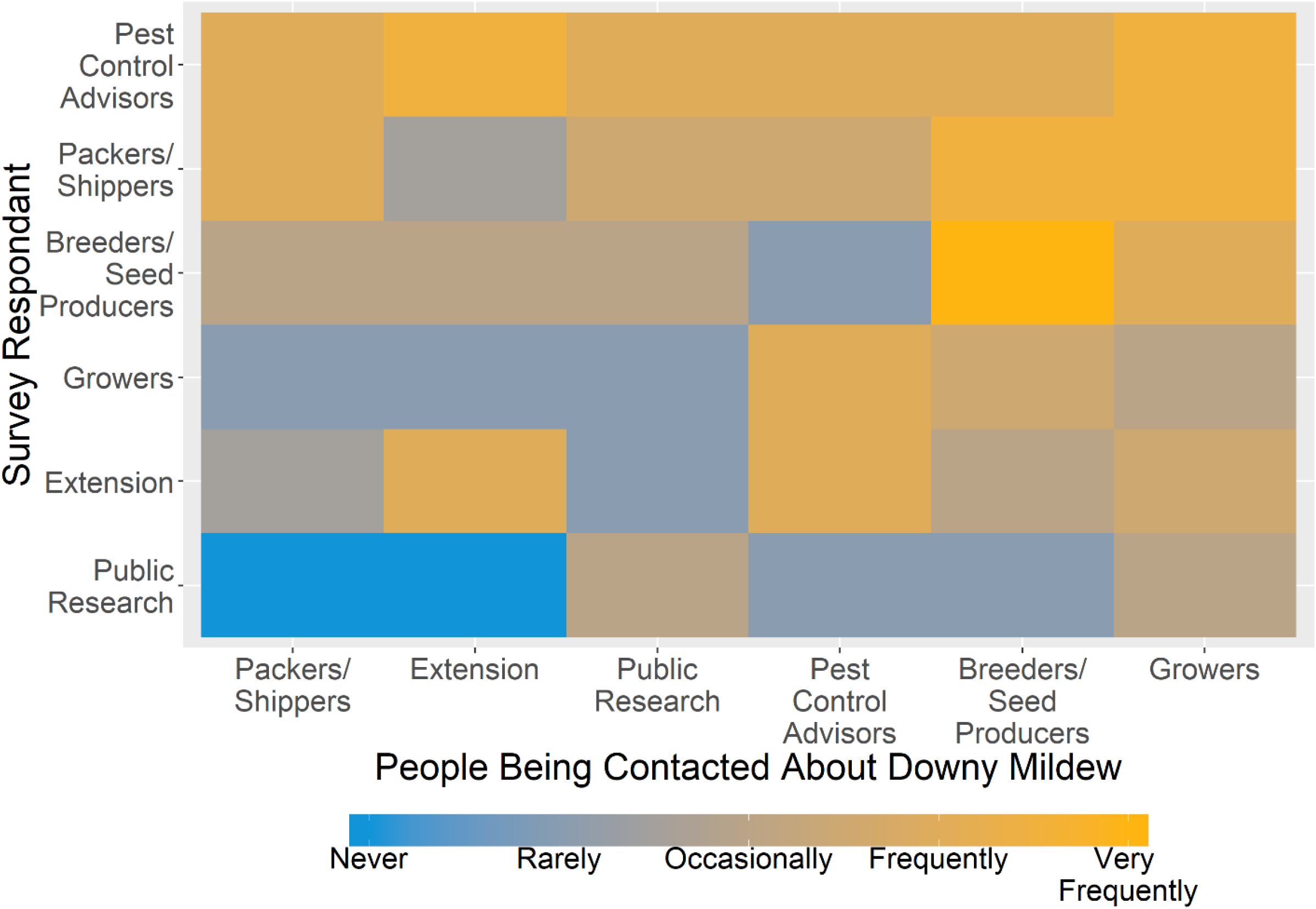
Communication matrix of people involved in spinach production. Colored boxes represent the median value of the frequency of communication between survey participant groups.

Disking diseased fields was perceived to have mixed efficacy by producers. The obligate biotrophic nature of *P. effusa* makes it difficult to impossible for the pathogen to reproduce on dead or dying plants. However, the recent discovery of durable oospore structures on seeds and in leaves of plants in production areas suggests that the pathogen maybe able to persist even through extreme management practices like field disking or plowing.

Roguing out diseased plants and the use of organic pesticides were perceived broadly to have low efficacy. Although roguing has been used as an effective management strategy for some diseases, this management is mostly limited to diseases that have short dispersal kernels. In addition, the nature of spinach production would require the regular hiring of field crews to remove diseased plants, which may be cost-prohibitive for many growers.

Despite these limitations, some managers have used field crews to rogue diseased plants in the past. Organic pesticides are available for spinach downy mildew control, however their efficacy is variable across sites, and can rarely be used to achieve economic levels of disease control.

## Demographics of Participants

The sum total of acreage managed by respondents was 33828 (n=23). This represents just over half of the 66150 acres of spinach planted in 2017 in the USA (USDA NASS 2017). It is possible (and even probable) that there was overlap in the acreage that different respondents claim responsibility for. Different respondent groups (e.g. – PCAs and growers) naturally overlap on the same acreage, and both play critical roles in production and disease management.

**Figure 2:**
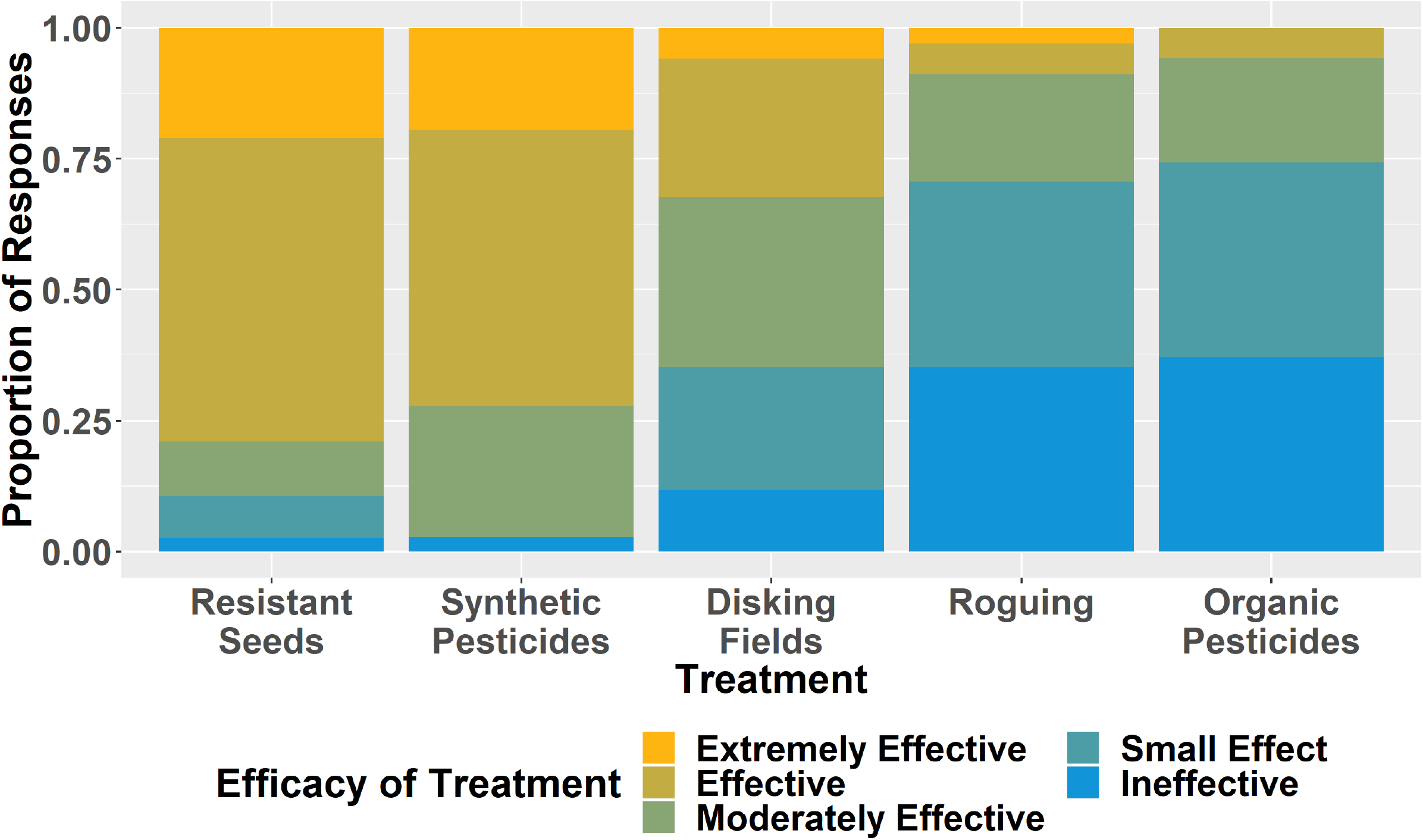
Perceived efficacy of different spinach downy mildew management practices by different respondent groups.

Only 27% of respondents (n=26) suggested that they regrow their spinach crops after harvesting. The practice helps to recoup some of the costs associated with expensive seeds (which can cost as much as $2000 per acre) but regrown crops may be more at risk for developing spinach downy mildew disease. Regrown crops are in the field for longer, and low levels of disease in the initial crop can rapidly expand to epidemic proportions in the secondary crop. Those that practice secondary harvests do so on average 47% (standard deviation 39%) of the time.

Respondents suggested that they see disease in approximately 19.7% of their fields. This proportion is lower than those found in other studies. Choudhury et al. (2016) found that 31.7% of fields surveyed for spinach downy mildew had disease incidence greater than 0.1%. Our participant’s estimate for disease affected fields may be lower because they do not count fields that had only small patches of disease, or that they were unable to detect disease due to insufficient surveillance.

Of the respondents who gave their age range (n=42), 47% responded that they were 55 years or older. Years of experience with spinach production for respondents (n=38) was 18.8 years on average (sd = 11). Of the respondents (n=41) who gave their education level, 93% had a college degree or an advanced degree.

## Conclusions and Future Prospects

Overall, the communication between groups is variable, with some groups communicating more regularly. PCAs seem to communicate frequently with nearly all participants in spinach production and may be good resources for finding out the current status of epidemics in the field and best management practices for managing disease. Most participants are contacting growers, suggesting strong involvement with the primary stakeholders of the industry. Increasing the communication between public researchers and extension agents may help to translate more basic research into practical management practices as well as help to shape future research goals.

Most people involved in spinach production are confident in the efficacy of resistant seeds and synthetic pesticides. These practices are the current standard managements for organic and conventional growers (respectively) and will hopefully remain viable into the future. The perceived efficacy of organic pesticides and roguing of diseased plants is low, suggesting that few participants place any amount of confidence in these practices. Increasing the number of publicly available research trials on both of these management practices may help to suggest if they are efficacious and improve adoption if they are.

Although relatively few growers regrow their crop, these plants may play a crucial role in disease dynamics as they may host disease at low levels, allowing it to extend its time in the production. The number of fields that growers estimate are diseased is lower than previously estimated, but still a sizable proportion of total production. Many of the participants in production are well educated and have years of experience in spinach production, suggesting that they know the system well and can adapt new management practices. The age range of participants is relatively top heavy, which is similar to agriculture as a whole. As more aging growers leave the industry and fewer young growers take their place, the industry may need to adapt to attract younger growers. The costs and uncertainty associated with managing spinach downy mildew may be preventing some young growers from entering the field.

In the future, surveys can be conducted to explore producer’s willingness to pay and adopt alternative management practices, like resistant seeds developed with CRISPR technology (Kock et al. 2019), spore trapping (Choudhury et al. 2016a), mixes of resistant and susceptible varieties (Choudhury et al. 2016b), oospore seed thresholds (Choudhury et al. 2017a), and organic pesticides (Choudhury et al. 2015a; b; Choudhury et al. 2017b). These studies could be conducted online or in person. In person surveys may help to resolve some of the ambiguities of the current study and help to improve response rates. As spinach downy mildew continues to pose a severe threat to spinach production, all areas of the spinach producing peoples need to adapt to adopt new management strategies.

**Figure 3:**
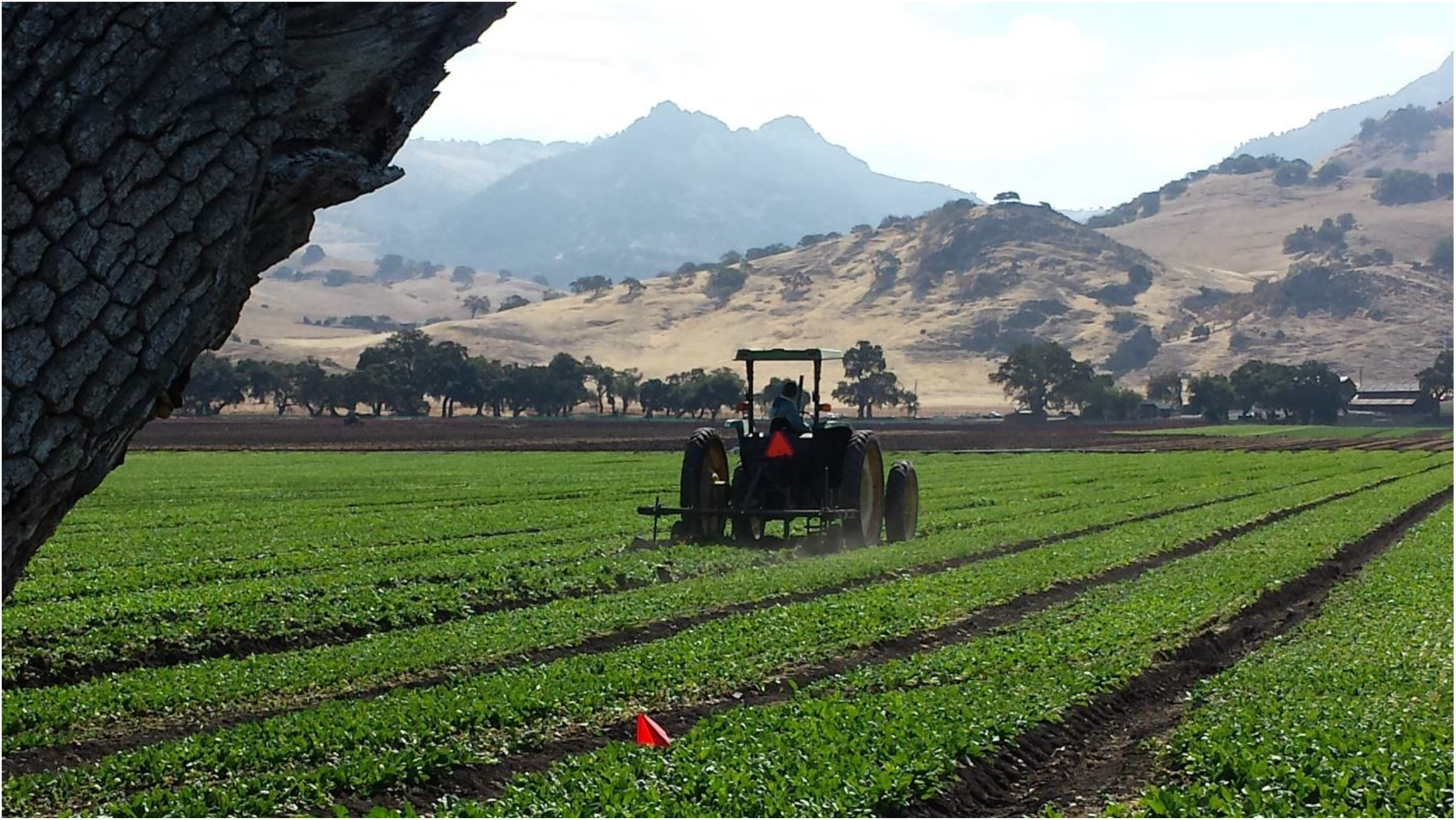
Mature spinach field affected by spinach downy mildew being disked into the ground by grower.

## Supporting information

Survey instrument

## Acknowledgements

We would like to thank the help of the California Leafy Greens Research Board who funded this project. We would especially like to thank Ms. Mary Ziske for help with coordinating communication with participants, and Mr Steven Koike for many helpful suggestions. We would also like to thank Drs. Mark Lubbell and Michael Levy for technical assistance and helpful comments in developing the survey questions.

