## Supplementary material for "Spinach Growers Value Host Resistance and Synthetic Fungicides in the Fight against Downy Mildew": Survey instrument

### Default Question Block

**About this study**
This survey is part of a research study of spinach production administered by UC Davis. Participation in this survey is voluntary. If you decide to complete this survey, it will take approximately 5 minutes of your time.
**All data from this survey will be kept anonymous. Your privacy is our priority.**
Thank you in advance.

If you have any questions, please contact:
Dr. Neil McRoberts, Associate Professor of Plant Pathology
530-752-3248


Which of the following best describes your primary role in spinach production?

- Grower
- Pest Control Advisor
- Breeder / Seed Producer
- Extension
- Public Research
- Packing / Shipping

Other (please specify)

### How frequently do you communicate with others in the industry for information on spinach downy mildew?

| | Never | Rarely | Occasionally | Frequently | Very Frequently |
|---|---|---|---|---|---|
| Growers | | | | | |
| Pest Control Advisors | | | | | |
| Breeders / Seed Producers | | | | | |
| Extension | | | | | |
| Public Research | | | | | |
| Packers and Shippers | | | | | |
| Other (please list) | | | | | |

### Please rank efficacy of spinach downy mildew control strategies

| | Ineffective | Small Effect | Moderately Effective | Effective | Extremely effective |
|---|---|---|---|---|---|
| Resistant seeds | | | | | |
| Synthetic Pesticides | | | | | |
| Organic Pesticides | | | | | |
| Disking of fields after harvest | | | | | |
| Use of field crew to remove infected plants | | | | | |
| Other (please list) | | | | | |

What percent of your spinach fields last year were affected by downy mildew?

What percent of your spinach fields last year were regrown after clipping?

How many acres of spinach do you typically manage in a year?

What is your age?

| | Younger than 25 | 25-34 | 35-44 | 45-54 | 55-64 | Older than 65 |
|---|---|---|---|---|---|---|
| Age | ○ | ○ | ○ | ○ | ○ | ○ |

What level of formal education have you completed?

| | High School, No Degree | High School, Degree | College, No Degree | College, Degree | Advanced Degree (e.g. - M.S.) |
|---|---|---|---|---|---|
| Formal Education | ○ | ○ | ○ | ○ | ○ |

How many years of experience do you have with spinach production?

Survey Powered By Qualtrics
